# dbGIST: An LLM-Assisted Multi-Omics Resource for Target Exploration and Cross-Dataset Validation in Gastrointestinal Stromal Tumors

**DOI:** 10.64898/2026.05.22.727292

**Authors:** Zhao Sun, Qian Zhao, Jie-Han Li, Jia-Jing Li, Hao Liu, Yan-Xu Guo, Ya-Dong Tang, Fei Yang, Xin Liu, San-Fei Peng, Wunan Mi, Ge Zhang, Zhen Zhang, Ming-Liang Yuan, Guang-Hui Li, Yun-Fei Wang, Chao Liu, Sheng-Lei Li, Jing-Hua Yang, Yang Fu

## Abstract

Gastrointestinal stromal tumors (GISTs) are the most common mesenchymal neoplasms of the gastrointestinal tract, yet GIST-specific omics evidence remains scattered across small cohorts and is not represented as a dedicated disease project in major cancer genomics resources, limiting reproducible target exploration. Here, we present dbGIST (https://www.dbgist.com), a dedicated GIST-focused multi-omics resource built to make dispersed GIST evidence searchable, analyzable, and reusable. dbGIST harmonizes data from 37 centers and 1,991 samples, including pathologically verified in-house cohorts, across genomics, bulk transcriptomics, proteomics, phosphoproteomics, and single-cell transcriptomics, and couples these data with curated clinical annotations covering survival, mutation status, risk stratification, metastasis or recurrence, mitotic index, tumor site and size, and imatinib response. The platform supports cohort-level molecular-clinical association, survival, enrichment, immune-infiltration, drug-sensitivity, and single-cell analyses through interactive visualizations, downloadable source data, and public APIs for programmatic access to reusable analysis outputs and visualization-ready data. An optional LLM-assisted interface helps users navigate analyses and interpret outputs. Using MCM7 as a case study, dbGIST linked a resource-derived candidate to survival, risk features, metastatic or recurrent disease, imatinib-response phenotypes, proliferative cell states, and in vitro GIST-cell behavior. dbGIST therefore provides a traceable and interoperable resource for target exploration and precision oncology research in GIST.

## 1. INTRODUCTION

Gastrointestinal stromal tumors (GISTs) are the most common mesenchymal neoplasms of the gastrointestinal tract and are characterized by substantial clinical and molecular heterogeneity [1]. Activating alterations in KIT and PDGFRA have established GIST as a paradigmatic disease for targeted therapy [2, 3]. Nevertheless, primary and secondary resistance to imatinib, metastatic progression, and biologically distinct non-canonical subgroups continue to complicate disease management and therapeutic target identification [4–10]. These challenges underscore the need for systematic resources that enable disease-focused interrogation of candidate genes, pathways, and clinically relevant vulnerabilities in GIST.

Multi-omics profiling has transformed target discovery in cancer [11]. However, GIST remains underrepresented in broad cancer reference resources such as The Cancer Genome Atlas (TCGA) and the International Cancer Genome Consortium (ICGC) [12, 13]. Consequently, widely used downstream platforms built on these compendia, including GEPIA2 and TIMER2.0, cannot directly support GIST-specific interrogation [14, 15]. Meanwhile, independent studies have generated valuable GIST datasets spanning genomics, transcriptomics, proteomics, phosphoproteomics, and single-cell transcriptomics, but these resources remain dispersed across repositories, generated by different platforms, and accompanied by uneven clinical annotation [16–18].

Beyond data fragmentation, practical barriers further limit reuse of existing GIST omics data: cross-modal comparison requires substantial manual harmonization, disease-relevant clinical variables are inconsistently organized, and many analytical workflows remain difficult to access for bench scientists and clinicians [19–21]. What the field lacks is therefore not merely another dataset collection, but a practical disease-focused resource for target exploration and cross-dataset validation. At the same time, recent advances in large language models (LLMs) create an opportunity to make such resources more interactive and accessible [22, 23]. For non-bioinformatic users, an LLM-powered conversational layer can provide guided platform navigation, user-oriented question answering, module-specific instructions, and interpretation of analytical outputs generated by the platform, thereby substantially improving usability [24, 25].

To address these needs, we developed dbGIST (https://www.dbgist.com), a disease-focused multi-omics resource that integrates dispersed public and three in-house GIST datasets into a unified analytical platform. dbGIST incorporates genomics, transcriptomics, proteomics, phosphoproteomics, and single-cell transcriptomics together with curated clinical annotations, and provides modules for clinical association analysis, survival analysis, functional enrichment, immune-related analysis, drug-response-related exploration, and cell-type-level interrogation. In addition, dbGIST includes an LLM-powered interactive interface that helps users understand how to use the platform, navigate different analytical modules, and interpret the resulting multi-omics outputs [26, 27]. We therefore designed dbGIST not only as a repository, but as an LLM-enhanced practical resource for target exploration and cross-dataset validation in GIST. To demonstrate this utility, we used MCM7 as a representative case and examined its behavior across cohorts, omics layers, and clinically relevant phenotypes, followed by experimental validation in vitro.

## 2. RESULTS

### 2.1. Overview of dbGIST

#### 2.1.1 Data and sample type

The dbGIST resource integrates a GIST-specific multi-omics collection spanning genomics, transcriptomics, proteomics, phosphoproteomics, and single-cell transcriptomics from 37 centers and 1,991 samples, including three in-house cohorts (Fig. 1A and 1B). Rather than treating these data as a single pooled matrix, dbGIST organizes them as modality-aware, cohort-level resources with explicit provenance. Public datasets were screened for GIST eligibility, diagnostic support, data type, platform, and clinical annotation availability; the resulting cohort manifests, and inclusion/exclusion records are provided in Supplementary Tables S1-S3. Quantitative outputs underlying key figures are provided as source-data tables where applicable. Transcriptomic and single-cell datasets were reprocessed from raw files where usable raw data were available, whereas published processed matrices were retained for proteomic/phosphoproteomic studies and other data types where raw reprocessing was not feasible or not applicable. This structure allows users to compare findings across cohorts while preserving the measurement scale and limitations of each source dataset.

**Figure 1:**
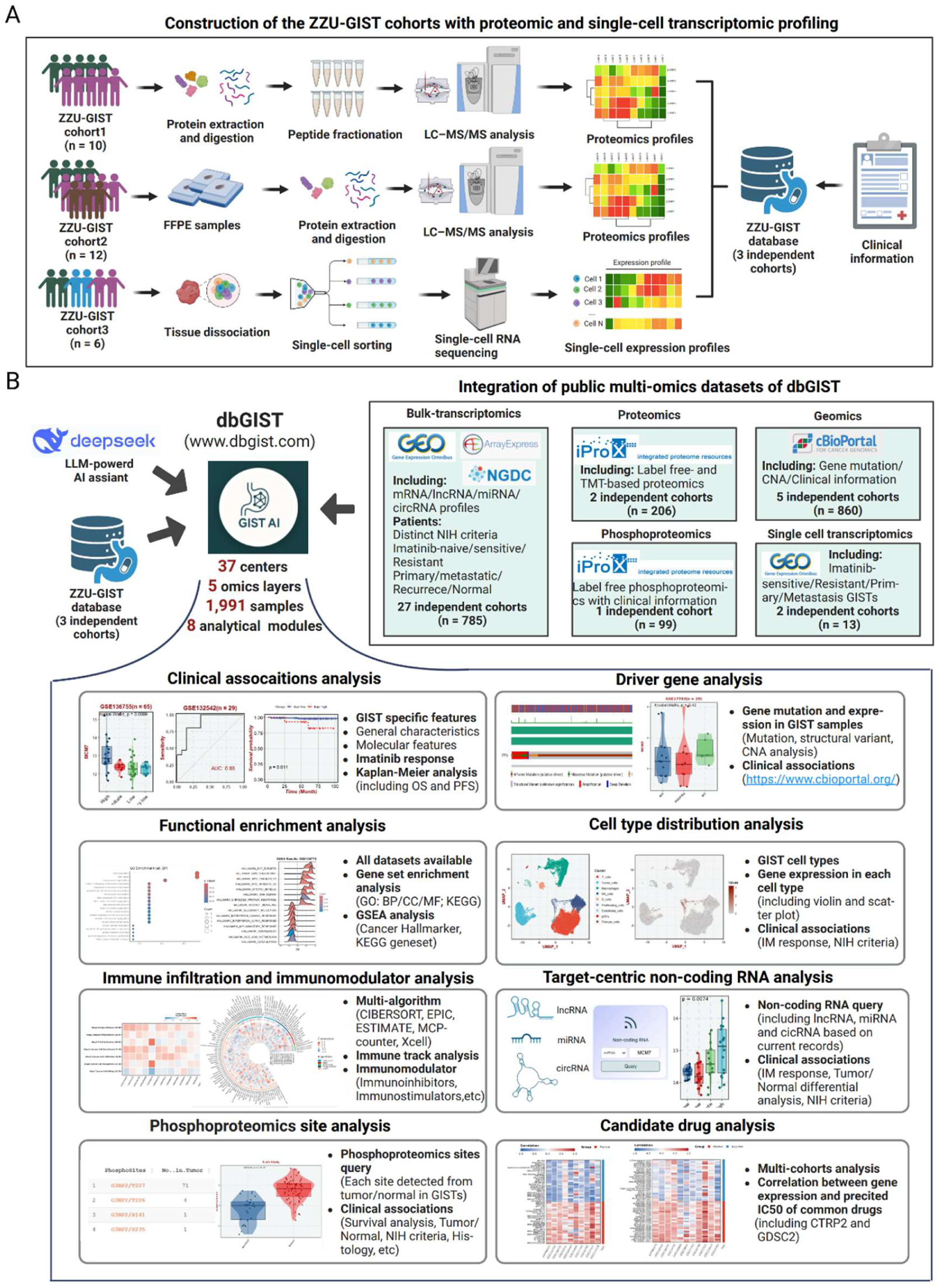
Overview of dbGIST as a GIST-focused multi-omics resource. (A) Sample processing of in-house GIST proteomics and single-cell transcriptomics. (B) Public and in-house GIST multi-omics data integration and main dbGIST analytical modules.

The included biospecimens and data objects cover surgical and biopsy-derived GIST specimens, tumor/normal contrasts where available, imatinib-sensitive and resistant settings, metastatic or recurrent disease, and multiple anatomic sites. Clinical variables were manually curated into controlled labels for common GIST-relevant phenotypes, including NIH risk, mutation status, metastatic status, mitotic index, tumor size, imatinib response, and survival endpoints. The three in-house ZZU-GIST cohorts add pathologically confirmed proteomic and single-cell transcriptomic data to the public collection, providing both a resource-building component and an independent context for cross-dataset validation (Fig. 1B).

#### 2.1.2 Home page with LLM-powered chatbot

dbGIST provides a web interface that combines curated multi-omics modules with an optional LLM-assisted interaction layer. The homepage includes a GIST-focused knowledge query interface and optional gene-centric literature support, including external literature-retrieval resources such as OpenScholar, to help users inspect published evidence for queried genes and navigate the resource (Fig. 2A; Supplementary Fig. S1A-C). This function is provided only for contextual exploration and user support; it does not contribute to dbGIST cohort-level statistical analyses or numerical results, which are generated by the platform’s predefined analytical modules.

**Figure 2:**
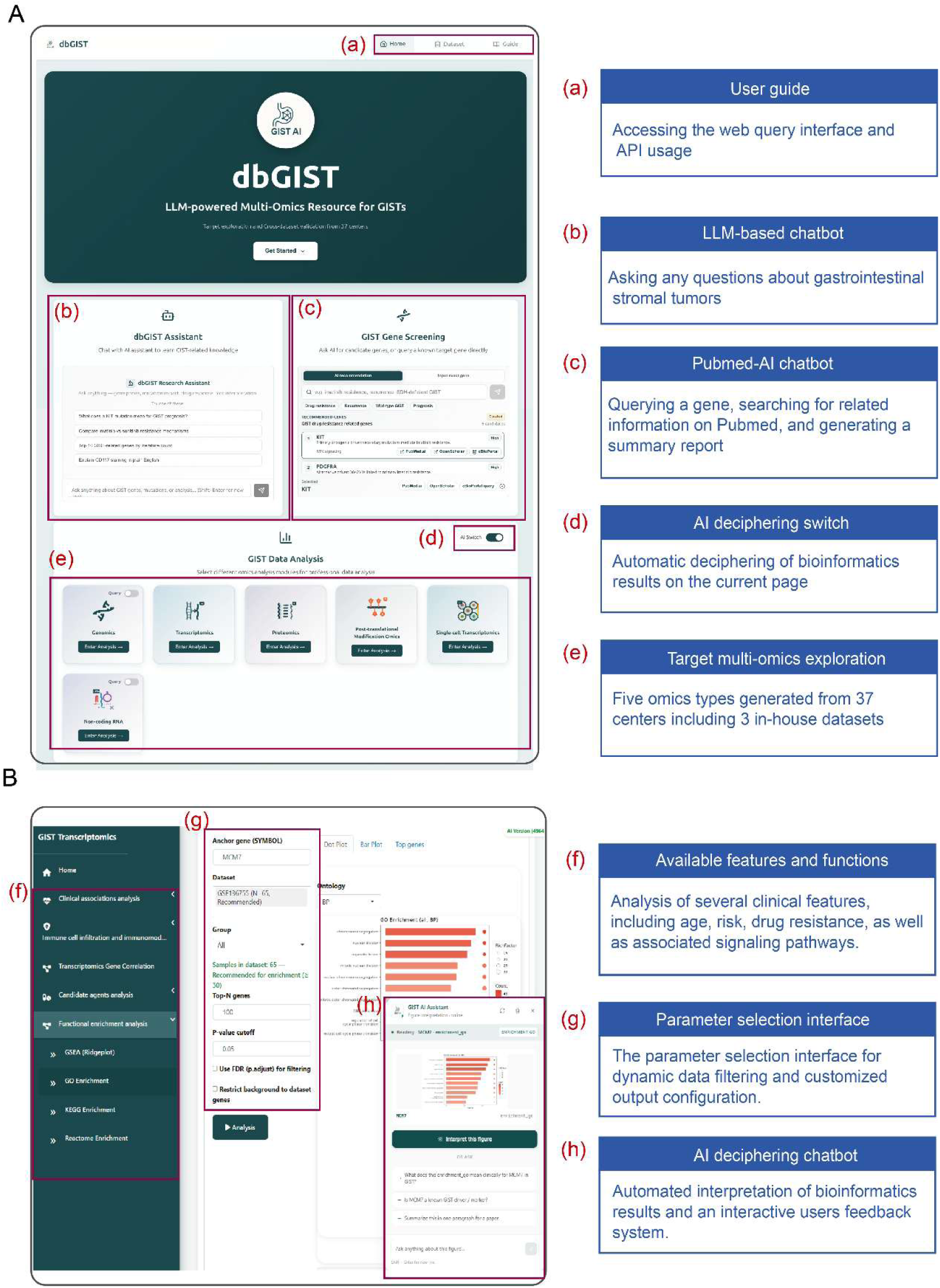
Overview of the dbGIST interface and analytical framework. (A) Homepage and core functional modules of dbGIST. (a) Top navigation bar providing access to Home, Dataset, and Guide pages for platform navigation and user support. (b) LLM-assisted GIST knowledge chatbot, enabling natural-language queries for biological questions and research guidance. (c) GIST gene screening module for candidate gene querying and target exploration. (d) AI interpretation switch that enables automatic explanation of bioinformatics results on the current page. (e) Entry points for multi-omics data analysis, including genomics, transcriptomics, proteomics, phosphoproteomics, single-cell transcriptomics, and non-coding RNA modules. (B) Representative omics analysis page (transcriptomics module). (f) Left-side navigation panel listing available analytical modules, including clinical association analysis, immune infiltration, gene correlation, and functional enrichment. (g) Parameter configuration interface allowing dataset selection, grouping strategy, and statistical threshold customization for dynamic analysis. (h) AI-assisted interpretation panel that provides automated explanation of analytical results and supports interactive user feedback.

Although dbGIST integrates five primary omics layers, its analytical capabilities are organized into six omics-focused sub-pages, with non-coding RNA evidence presented as a separate evidence-aggregation page. The optional LLM-powered interpretation module can be enabled before an omics-specific analysis to support plain-language explanation of figures, module-specific guidance, and follow-up questions (Fig. 2B). This layer is intended to improve accessibility, but it does not alter preprocessing, statistical testing, or returned numerical results. In parallel, the platform provides downloadable visualization outputs and structured result data, and public API endpoints expose selected analysis results and visualization-ready data for external workflow integration.

#### 2.1.3 Omics data analysis page

The dbGIST platform provides eight core analytical modules that enable systematic and interactive exploration of GIST multi-omics data. As summarized in Fig. 1B, these modules cover clinical association analysis, driver-gene analysis, functional enrichment, cell-type distribution, immune infiltration, non-coding RNA evidence, phosphoprotein-site analysis, and candidate-drug exploration. Each module operates on curated, precomputed data objects and returns cohort-aware outputs rather than LLM-generated statistics. Users initiate analyses by submitting a gene, after which dbGIST checks data availability and executes the corresponding workflow. Results include publication-ready figures and structured result tables, and selected derived outputs can also be queried through public API endpoints.

### 2.2. Exploring the multi-omics functions of dbGIST using MCM7 as an illustrative example

To demonstrate how dbGIST can support resource-derived hypothesis generation in gastrointestinal stromal tumor (GIST) research, we selected MCM7 (Minichromosome Maintenance Complex Component 7), a key regulator of DNA replication initiation and cell cycle progression, as a representative query gene [28]. The literature-oriented interface, including external retrieval resources such as OpenScholar, was used only to provide background context for this illustrative query. We then used MCM7 as an illustrative candidate to explore how dbGIST connects transcriptomic expression, clinical associations, single-cell distribution, functional enrichment, immune context, drug-response prediction, and experimental follow-up. No recurrent genomic alteration or quantified phosphoproteomic event for MCM7 was detected in the current GIST resource; therefore, the case study focuses on data layers in which MCM7 was measured and informative, while the same modules remain available for genes covered by genomic or phosphoproteomic datasets.

#### 2.2.1 Clinical associations

The dbGIST platform enables systematic investigation of associations between molecular features and clinically relevant parameters across integrated omics modalities. Clinical variables were manually curated and harmonized where the source studies allowed, including imatinib treatment response, NIH risk stratification, metastatic status, driver mutation profiles, mitotic index, tumor size, tumor location, age, sex, and survival endpoints. An overview of the primary clinical variables available for each omics dataset is provided in Fig. 3A. When a user initiates a clinical correlation analysis by entering a gene of interest, dbGIST first checks whether the queried feature is available in each applicable cohort and then performs modality-specific association analyses within those cohorts, returning the results for interactive exploration.

**Figure 3:**
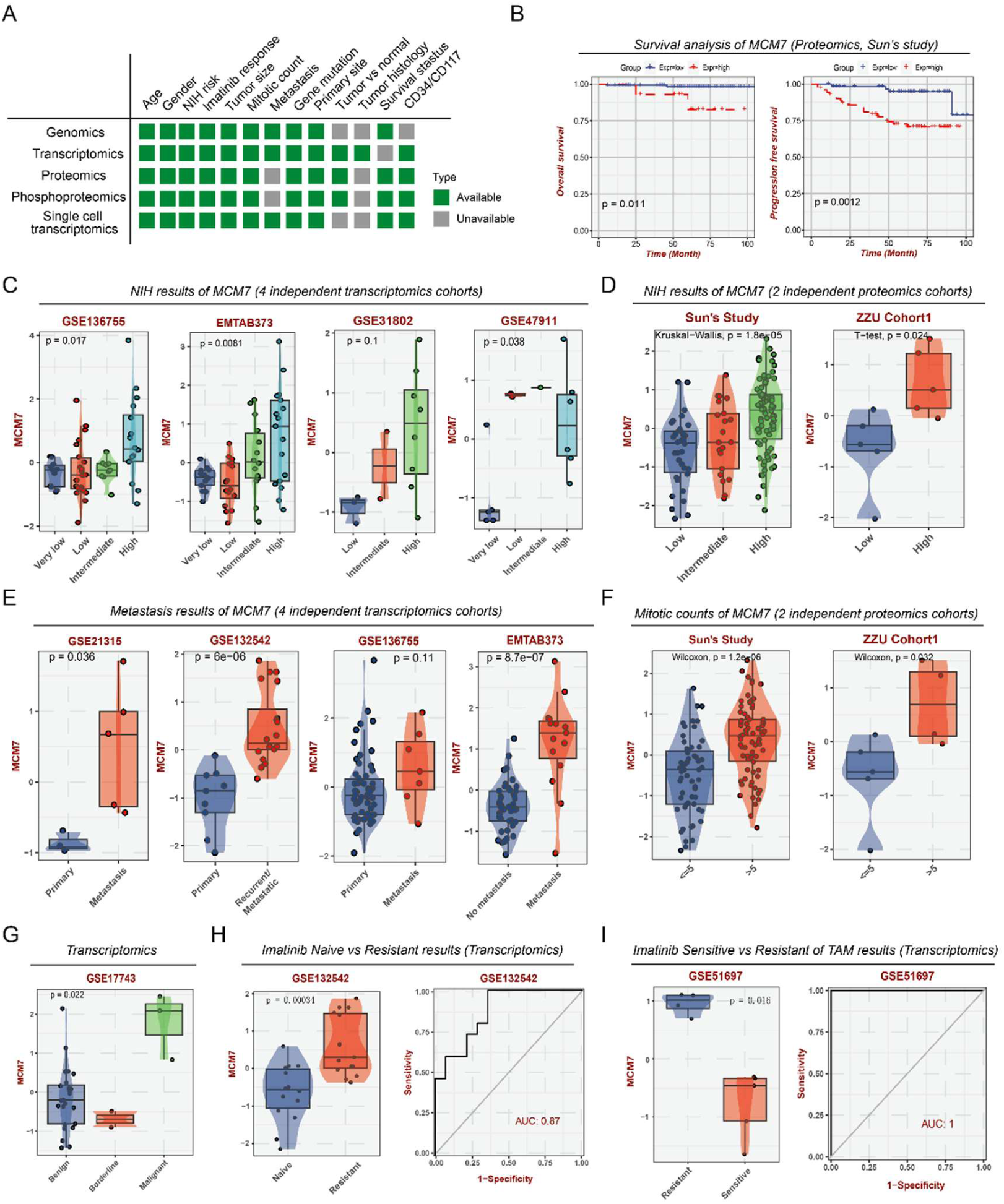
Clinical associations of MCM7 based on integrated omics data. (A) Summary of clinical annotations available for each omics layer. (B) Kaplan-Meier curves showing overall survival (OS) and progression-free survival (PFS) in the proteomics cohort stratified by MCM7 expression. (C) MCM7 expression across NIH risk groups in four independent transcriptomic cohorts. (D) MCM7 expression across NIH risk groups in two proteomic cohorts. (E) MCM7 expression across primary, metastatic, recurrent/metastatic, and non-metastatic comparison groups in four transcriptomic cohorts. (F) Association between MCM7 expression and mitotic count in two proteomic cohorts. (G) MCM7 expression across histologic categories in a transcriptomic cohort. (H) MCM7 expression by imatinib response and ROC analysis for distinguishing imatinib-resistant from imatinib-naive cases in GSE132542. (I) MCM7 expression by imatinib response and ROC analysis in tumor-associated macrophages from GSE51697. Centre line: median; box bounds: 25th and 75th percentiles.

As shown in Fig. 3B, proteomic data revealed that high MCM7 expression was significantly associated with poorer overall survival (OS) and progression-free survival (PFS) (p = 0.011 and p = 0.0012, respectively). Transcriptomic and proteomic analyses further linked MCM7 expression to NIH risk classification (Fig. 3C-D). Elevated MCM7 levels in high-risk patients were observed in three independent transcriptomic cohorts (GSE136755: p = 0.017; E-MTAB-373: p = 0.0081; GSE47911: p = 0.038), a finding corroborated in the in-house ZZU-Cohort1(p = 0.024). MCM7 was also higher in metastatic or recurrent tumors than in non-metastatic cases across three transcriptomic datasets (GSE21315: p = 0.036; GSE132542: p = 6e-06; E-MTAB-373: p = 8.7e-07). Mitotic index showed a positive association with MCM7 expression (Fig. 3F; p = 1.2e-06 and p = 0.032 in two cohorts), and MCM7 was upregulated in malignant histologic subtypes compared with benign GIST samples (p = 0.022). In imatinib-response analyses, MCM7 expression was elevated in resistant cases (p = 0.00034) and showed discriminatory performance for resistance (AUC = 0.87; Fig. 3H). In a transcriptomic dataset derived from tumor-associated macrophages isolated from GIST tissues, MCM7 yielded an AUC of 1.0 in this cohort (Fig. 3I). Together, these cohort-level findings nominate MCM7 as a clinically relevant, resource-derived candidate associated with risk, survival, metastatic status, and treatment-response phenotypes.

#### 2.2.2 Cell type distributions

The dbGIST platform enables integrated exploration of single-cell transcriptomic data across multiple independent cohorts. It incorporates three curated single-cell datasets, including two public datasets (GSE162115 and GSE254762) and one in-house ZZU-GIST cohort, processed with standardized quality control, normalization, dimensionality reduction, and cell-type annotation steps (see Methods and Supplementary Methods). Users can query any gene of interest to retrieve its cell-type-specific expression prevalence and perform differential expression analysis across pre-annotated cell populations. Comparative expression analysis can also be performed within clinically stratified cell subsets, such as imatinib-resistant versus sensitive or high-risk versus low-risk groups, enabling cell-level assessment of gene expression patterns in clinical context.

Following standardized processing, the GSE162115, GSE254762, and ZZU-Cohort3 yielded 31,870, 55,429, and 60,437 high-quality cells, respectively. Additional before-QC, after-QC, no-Harmony, and Harmony assessment panels for all three cohorts are provided in Supplementary Figure S4A-L. The cellular composition and subtype distribution for each dataset are summarized in Fig. 4A, 4D, and 4G. MCM7 was consistently enriched in proliferative cell clusters across all three cohorts (Fig. 4B-C, E-F, H-I). These proliferative clusters included malignant cells and actively cycling T cells, both characterized by elevated proliferation markers such as MKI67. dbGIST further linked MCM7 expression in tumor cells to clinically relevant traits, including NIH risk grade, recurrence status, mitotic index, and imatinib resistance (Fig. 4J-M). These results demonstrate that the single-cell module can connect gene-level expression patterns with cell states and clinical annotations across cohorts.

**Figure 4:**
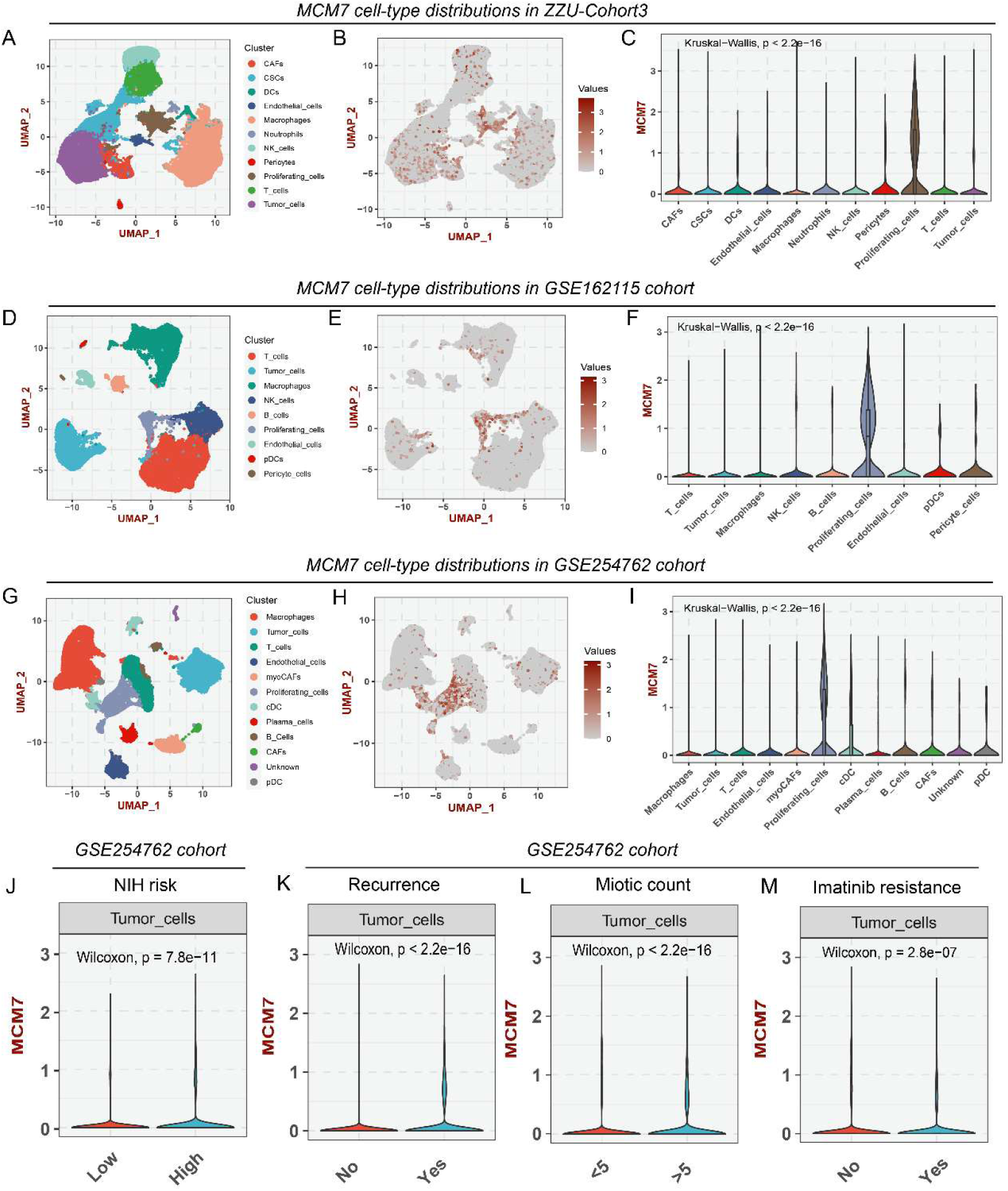
Single-cell distribution and tumor-cell associations of MCM7 in GIST. (A, D, G) UMAP visualization of annotated cell types in the ZZU-GIST, GSE162115, and GSE254762 cohorts, respectively. (B, E, H) UMAP feature plots showing MCM7 expression in each cohort. (C, F, I) Violin plots showing MCM7 expression across annotated cell types. (J-M) MCM7 expression in tumor cells stratified by NIH risk in GSE162115 and by recurrence, mitotic count, and imatinib resistance in GSE254762. Centre lines indicate medians where shown.

#### 2.2.3 Biological functional enrichment analysis

The dbGIST platform enables users to explore target-gene function through over-representation analysis and gene set enrichment analysis (GSEA), with the integrated workflow shown in Fig. 5A. We used this module to characterize the functional landscape associated with MCM7 expression in GIST. Interrogation of the transcriptomic dataset GSE136755 by Hallmark GSEA revealed enrichment of proliferation-associated pathways, including G2M checkpoint and mitotic spindle gene sets in samples with high MCM7 expression (Normalized Enrichment Score [NES] > 2.0, False Discovery Rate [FDR] < 0.001; Fig. 5B). Over-representation analysis of top MCM7-correlated genes further highlighted cell-cycle-related biological processes, including cell division and nuclear division (Fig. 5C), and network visualization illustrated an interconnected mitotic-regulation module (Fig. 5D).

**Figure 5:**
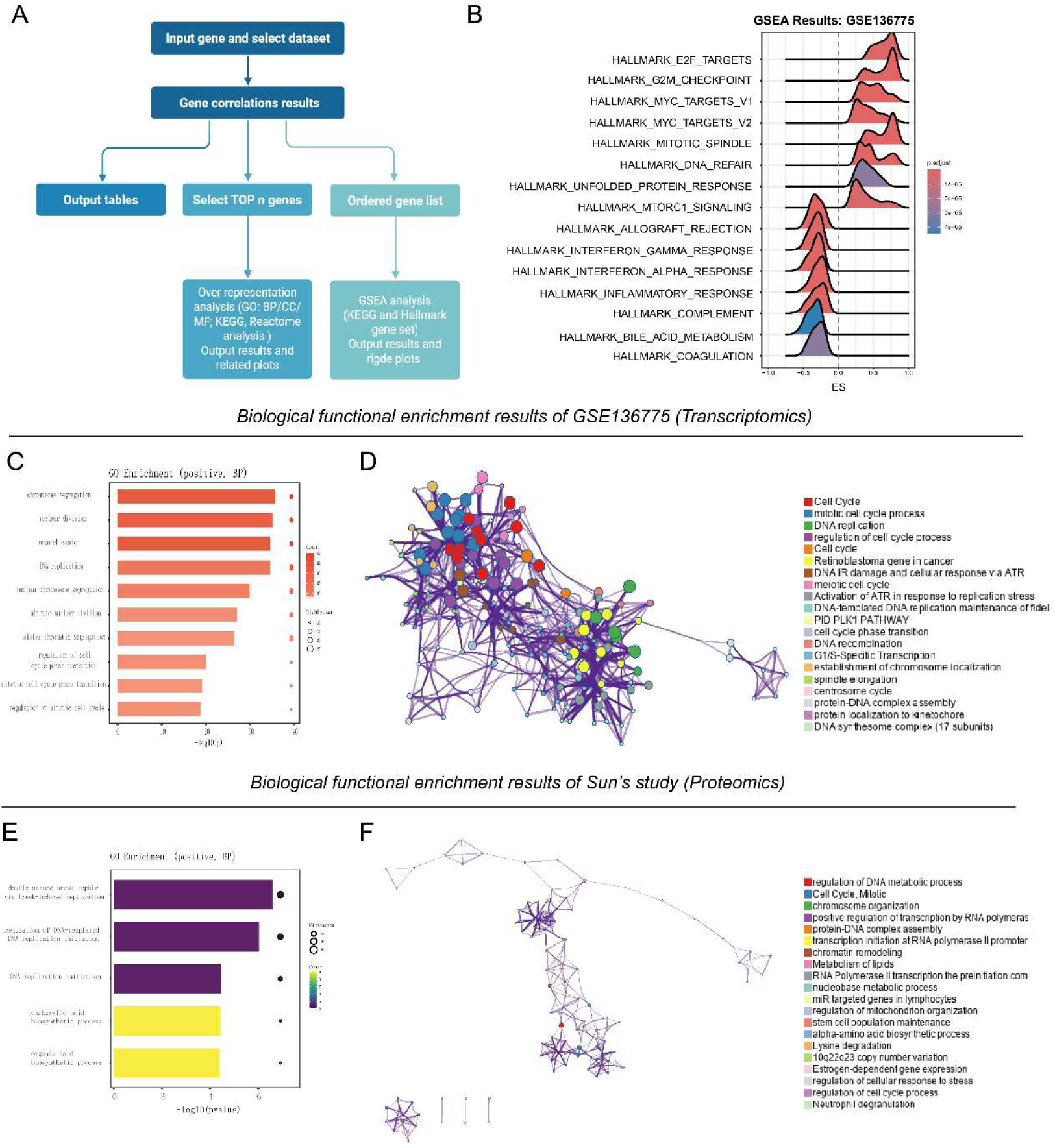
Biological pathway enrichment results of MCM7 in GIST. (A) Schematic workflow of the integrated bioinformatic analysis pipeline. (B) Gene Set Enrichment Analysis (GSEA) plot of the Hallmark gene set. (C) Bar graph from over-representation analysis (ORA) of the top MCM7-correlated genes. (D) Protein-Protein interaction network of the top MCM7-correlated genes. (E) Bar graph from over-representation analysis (ORA) of the top MCM7-correlated proteins. (F) Protein-Protein interaction network of the top MCM7-correlated proteins.

At the protein level, GO biological-process analysis of MCM7-correlated proteins identified DNA replication as the most significantly enriched pathway (Fig. 5E), reinforcing the transcriptomic findings. Network analysis further connected MCM7-associated proteins with cell-cycle and chromosome-organization processes (Fig. 5F). Together, these transcriptomic and proteomic analyses support a reproducible proliferative signature associated with MCM7-high GIST samples, without requiring pooled inference across heterogeneous cohorts.

#### 2.2.4 Immune infiltration, immunomodulators, and candidate compounds

The dbGIST platform also enables systematic investigation of target molecules within the tumor immune microenvironment and candidate-drug context (Fig. 6A). Immune infiltration and tumor microenvironment features are estimated using complementary deconvolution frameworks, including xCell, CIBERSORT, and related tools [29, 30]. Drug-response analyses use pharmacogenomic prediction models trained on large public drug-screening resources, with derived results cached in the platform for rapid retrieval. Users can visualize cohort-level correlations, inspect scatterplots, and download structured result data for downstream interpretation or API-based workflow integration.

**Figure 6:**
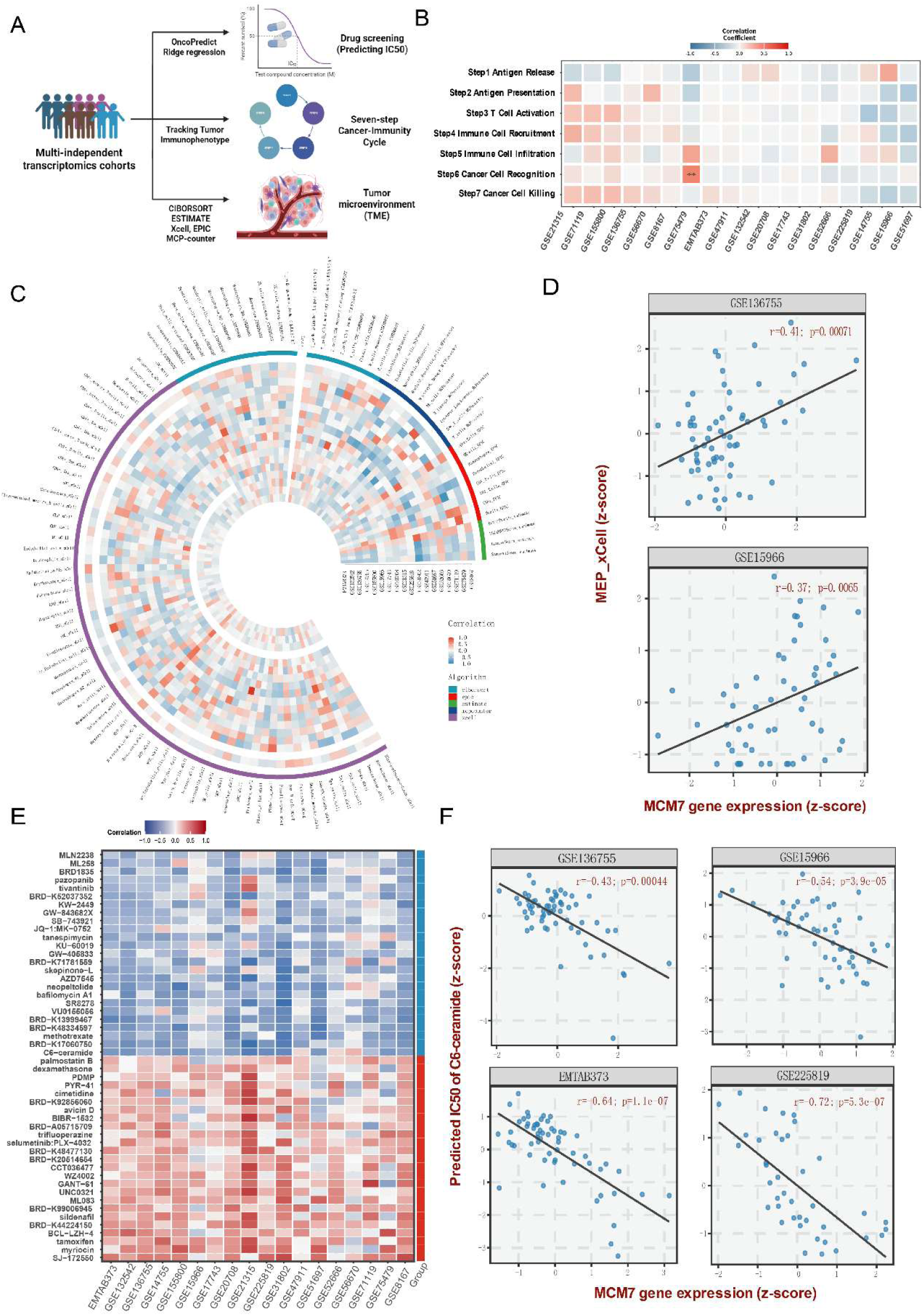
Immune infiltration and drug-sensitivity analyses of MCM7. (A) Flowchart of the integrated bioinformatic pipeline used for tumor immunophenotyping and in silico drug prediction across multiple GIST cohorts. (B) Heatmap showing Pearson correlation coefficients between MCM7 expression and activity of the seven-step cancer-immunity cycle. Red indicates positive correlation and blue indicates negative correlation. (C) Circos plot showing correlations between MCM7 expression and immune cell subtype abundance in the tumor microenvironment, as estimated by multiple deconvolution algorithms. (D) Scatter plots showing positive correlations between MCM7 and MEP (megakaryocyte-erythroid progenitor) cell abundance. (E) Heatmap of correlations between MCM7 expression and predicted drug sensitivity or resistance across analyzed cohorts. (F) Scatter plots showing significant negative correlations between MCM7 expression and predicted response to C6-ceramide.

As illustrated in Fig. 6B, analysis of MCM7 within the cancer-immunity cycle showed no strong association with antigen release or presentation steps. Tumor microenvironment deconvolution further indicated that higher MCM7 expression was inversely correlated with cytotoxic T-cell and natural-killer-cell infiltration (Fig. 6C). dbGIST also identified a positive correlation between MCM7 and MEP (megakaryocyte-erythroid progenitor) cell abundance estimated by deconvolution across independent cohorts (GSE136755: r = 0.41, p = 0.00071; GSE15966: r = 0.37, p = 0.0065), with cohort-level scatterplots available for inspection (Fig. 6D). In the drug-sensitivity module, C6-ceramide showed a consistent negative correlation with MCM7 expression across multiple datasets (GSE136755: r = -0.43, p = 0.00044; GSE15966: r = -0.54, p =3.9e-05; E-MTAB-373: r = -0.64, p = 1.1e-07; GSE225819: r = -0.72, p = 5.3e-07), nominating it as a candidate compound for further experimental evaluation in MCM7-high GIST contexts (Fig. 6F).

#### 2.2.5 Biological experimental validation of MCM7 in vitro GIST models

To evaluate whether the resource-derived MCM7 signal reflected a testable biological phenotype, we first assessed its relationship to proliferation markers within dbGIST. Across multiple independent patient cohorts, MCM7 showed a positive correlation with the established proliferation marker MKI67 (Ki-67) (Fig. 7A), supporting a link between MCM7 expression and proliferative activity. We then established MCM7-knockdown models in two GIST cell lines, GIST-T1 and GIST-882, using specific siRNAs. Western blot analysis confirmed efficient reduction of MCM7 protein in si-MCM7-transfected cells relative to controls (Fig. 7B, D), and quantitative real-time reverse-transcription PCR further verified reduced MCM7 mRNA expression (Fig. 7C, E).

**Figure 7:**
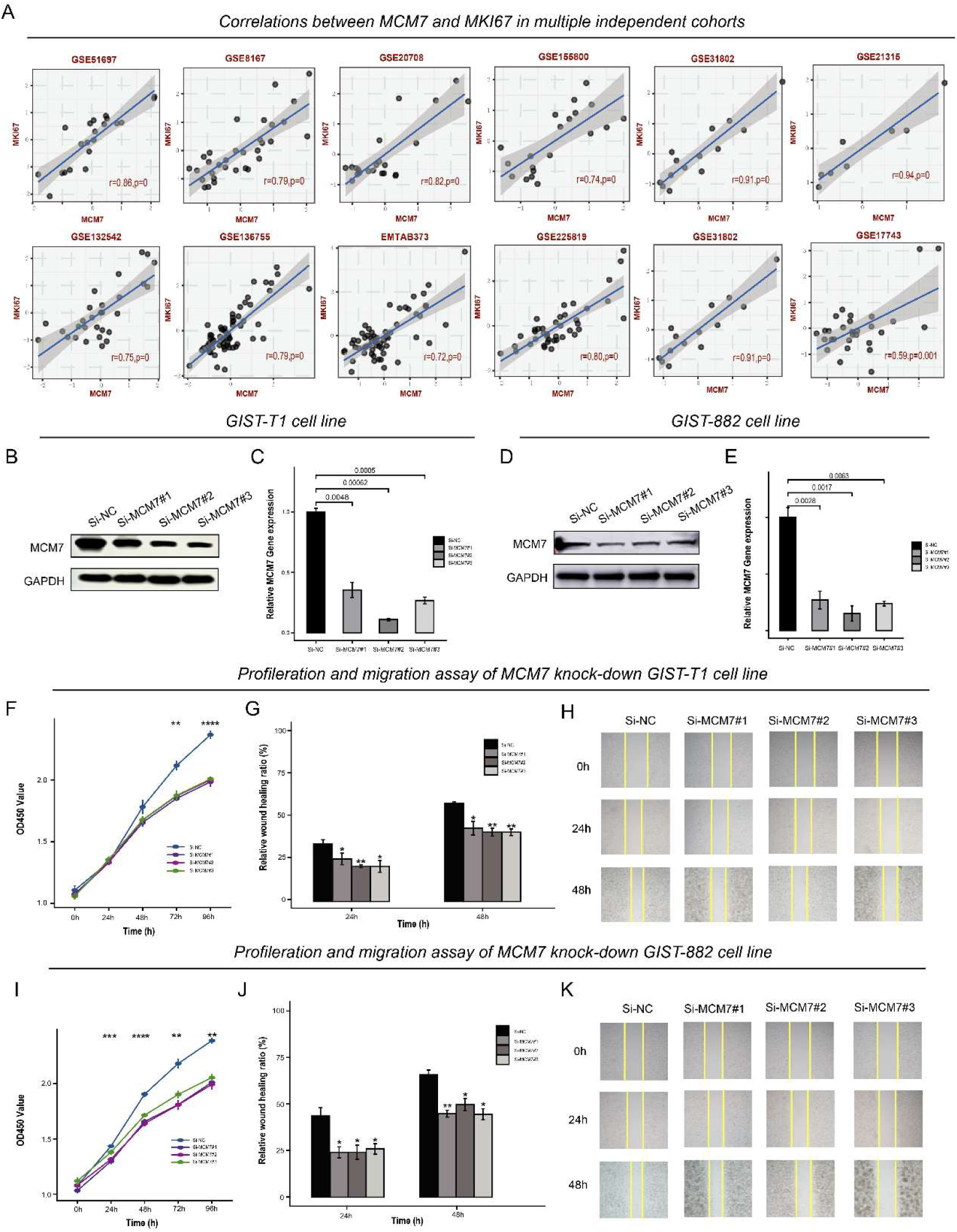
Experimental validation of MCM7 in in vitro GIST models. (A) Scatter plots showing correlations between MCM7 and MKI67 expression across independent GIST cohorts. (B, D) Western blot validation of siRNA-mediated MCM7 knockdown in GIST-T1 and GIST-882 cells. (C, E) qRT-PCR analysis of MCM7 mRNA expression after siRNA-mediated knockdown in GIST-T1 and GIST-882 cells. (F, I) CCK-8 proliferation assays after MCM7 knockdown in GIST-T1 and GIST-882 cells. (G, J) Quantification of wound-healing assays. (H, K) Representative wound-healing images at 0, 24, and 48 h. Statistical significance: *P < 0.05; **P < 0.01; ***P < 0.001.

Functional assays showed that MCM7 knockdown impaired the proliferative capacity of both GIST-T1 and GIST-882 cells (Fig. 7F, I). Wound-healing assays further indicated reduced migratory potential after MCM7 depletion (Fig. 7G, H, J, K). These experiments provide biological support for the dbGIST-derived prioritization of MCM7 and illustrate how the resource can generate experimentally testable target hypotheses.

## 3. DISCUSSION

In this study, we present dbGIST as a disease-focused multi-omics resource for gastrointestinal stromal tumors rather than a conventional data archive. By integrating datasets from 37 centers, 1,991 samples, three in-house GIST cohorts, and five omics layers, dbGIST addresses a long-standing gap created by the absence of a dedicated GIST cohort in broad cancer genomics consortia. Its value lies in combining curated molecular measurements, clinically meaningful annotations, interactive analytical modules, and accessible interpretation tools within a single GIST-specific research infrastructure.

A central contribution of dbGIST is the curation and harmonization framework underlying the resource. Public GIST omics datasets are heterogeneous in platform, file type, clinical annotation depth, and diagnostic metadata. We therefore verified GIST cohort eligibility by cross-referencing repository records, primary publications, and supplementary clinical files; recorded ICD or ICD-O codes when available; excluded ambiguous mixed tumor collections; and screened TCGA-related categories without identifying a dedicated GIST cohort. Where raw files were usable, transcriptomic and single-cell datasets were reprocessed with modality-appropriate workflows, whereas published processed matrices were retained for data types where raw reprocessing was not feasible or not applicable. Statistical analyses are performed within cohorts, with cross-cohort displays used for comparative interpretation rather than pooled inference. This design makes the scope and limits of harmonization explicit while preserving biological interpretability.

The three in-house ZZU-GIST cohorts further strengthen dbGIST by adding deeply profiled, pathologically confirmed GIST data to the public collection. These cases were confirmed by experienced pathologists using histomorphology and institutional immunohistochemical evaluation, including CD117/KIT and DOG1 where available, with clinicopathological information reviewed from pathology reports and medical records. Importantly, the resource is designed for reuse beyond the web interface. dbGIST provides public API access to analysis results and the result-associated data used for visualization, enabling external tools to query GIST-specific evidence programmatically and incorporate dbGIST outputs into independent analytical workflows.

The MCM7 case study illustrates how dbGIST can support resource-derived hypothesis generation and validation. Across transcriptomic, proteomic, clinical, and single-cell contexts, MCM7 was associated with high-risk features, imatinib response-related phenotypes, adverse survival, and proliferative cell states [28, 31]. Functional perturbation in GIST cell models further supported a role for MCM7 in tumor cell proliferation. We therefore view MCM7 not as an isolated single-gene endpoint, but as a representative example of how dbGIST can connect cross-dataset evidence, multi-omics context, and experimental follow-up for target exploration in GIST.

A cornerstone of dbGIST’s utility lies in its LLM-powered framework, which we have successfully implemented to fundamentally transforms the user experience by automating the interpretation of analytical results [26]. The conversational AI assistant fundamentally transforms the user experience by automating the interpretation of analytical results, for instance, explaining the clinical implications of a survival curve or the biological context of a pathway enrichment analysis in plain language[32]. More importantly, it provides real-time, dialogue-based troubleshooting, effectively guiding users with limited computational expertise through the analytical workflow. This significantly lowers the barrier for clinicians and wet-lab researchers, empowering them to directly mine the integrated dataset for insights without relying on specialized bioinformatics support. Beyond providing a set of tools, we have established a scalable and transferable framework for building intelligent, domain-specific research platforms (https://github.com/FuLab-ZhaoSun/dbGIST). This framework can potentially be adapted to other rare or understudied malignancies that similarly suffer from fragmented data resources and a lack of dedicated analytical services.

Several limitations should be considered. Public GIST datasets remain uneven in sample size, platform coverage, clinical variables, and available raw data, and phosphoproteomic coverage is still more limited than transcriptomic or single-cell coverage. The LLM layer is designed as an optional interpretation and navigation assistant; it does not modify the statistical backend and will require continued maintenance, monitoring, and literature updates. These literature-oriented AI functions, including external retrieval resources such as OpenScholar, are intended to facilitate user navigation and background exploration, but should not be interpreted as systematic reviews or substitutes for manual literature verification [27]. Future versions of dbGIST will expand cohort coverage, improve API documentation, incorporate emerging modalities such as spatial transcriptomics when available, and maintain the platform as a versioned, interoperable, community-oriented resource for GIST research.

## 4. METHODS

### 4.1 Data collection, cohort eligibility, and pre-processing

We systematically retrieved GIST-related multi-omics datasets from public repositories, including GEO, ArrayExpress, iProX, GSA-Human/NGDC, and cBioPortal, together with in-house cohorts generated at the First Affiliated Hospital of Zhengzhou University. Dataset inclusion required explicit study-level support for a GIST cohort, availability of quantifiable molecular measurements, and accessible clinical annotations. Cohort verification was performed by cross-referencing repository metadata, the primary publication, and supplementary clinical files where available. ICD or ICD-O coding was recorded when explicitly provided, but was not treated as a mandatory inclusion field because such registry-style variables are not consistently exposed in public omics repositories. Datasets labeled only as generic sarcoma, gastrointestinal tumor, or mixed mesenchymal neoplasm were excluded unless a GIST-specific subset could be unambiguously resolved. Detailed cohort manifests, inclusion or exclusion records, and diagnosis-verification notes are provided in Supplementary Tables S1-S3 and Supplementary Methods Section 1.

In-house ZZU-GIST specimens were obtained from surgically treated patients at the First Affiliated Hospital of Zhengzhou University under institutional ethics approval (Approval No., 2024-KY-1592-001 and 2024-KY-2222-02), with written informed consent obtained from all participants. All included cases were pathologically confirmed as gastrointestinal stromal tumors by experienced pathologists based on histomorphology and institutional immunohistochemical evaluation, including CD117 (KIT) and DOG1 where available. Clinicopathological information was reviewed from pathology reports and medical records, and KIT/PDGFRA mutation status was incorporated when available for molecular annotation and subgroup analysis.

Pre-processing was performed in a modality-aware manner rather than through a single merged pipeline. For transcriptomics, raw files were reprocessed with platform-specific workflows whenever usable raw data were available. Genomic cohorts were harmonized to a common mutation-annotation framework centered on the MSK-IMPACT convention. Published quantification matrices were retained for proteomics and phosphoproteomics, whereas single-cell datasets were processed individually with a standardized Seurat-based workflow before being incorporated into the platform as precomputed objects. As part of resource curation, we also screened TCGA soft tissue sarcoma and related disease categories for potential GIST cases, but no dedicated GIST cohort was identified; TCGA was therefore not used as a GIST sample source in dbGIST. Full modality-specific workflows are summarized in Supplementary Methods Section 1.

Clinical annotations were harmonized through a controlled vocabulary rather than by forcing all cohorts into one pooled analytic matrix. Imatinib response, NIH risk class, mutation category, metastatic status, and other recurrent variables were recoded into standardized labels wherever the source studies allowed. Importantly, inferential analyses in dbGIST are performed within individual datasets, and cross-cohort displays are used for comparative interpretation rather than for batch-corrected pooled inference. Gene-wise Z-score transformation is therefore used only for visual comparability across cohorts, whereas statistical testing remains cohort-specific. This analytical scope is detailed further in Supplementary Methods Section 1.

### 4.2 In-house ZZU-GIST proteomics

The in-house ZZU-GIST proteomics dataset comprised two independent cohorts with distinct sample-processing workflows: liquid nitrogen–frozen tissue specimens in ZZU-Cohort 1 and paraffin-embedded tissue section samples in ZZU-Cohort 2. For ZZU-Cohort 1, proteins were extracted using a RIPA-based workflow, digested with trypsin, fractionated by high-pH reversed-phase chromatography, and analyzed by LC–MS/MS on an EASY-nLC 1200 system coupled to a Q Exactive HF-X Orbitrap mass spectrometer. For ZZU-Cohort 2, deparaffinized tissue sections were subjected to SDS-based protein extraction, tryptic digestion, and LC–MS/MS analysis on an EASY-nLC 1200 system coupled to an Orbitrap Exploris 480 mass spectrometer in data-dependent acquisition mode.

Raw mass spectrometry data were processed using Proteome Discoverer with cohort-specific human reference protein databases and search parameters. False discovery rates were controlled at 1% according to the corresponding database-searching workflows. Protein expression matrices were generated from identified proteins and their label-free quantification intensity values for downstream analyses. Detailed sample-preparation, LC–MS/MS acquisition, database-searching, and quantification procedures are provided in Supplementary Methods Section 1.4.3.

### 4.3 In-house ZZU-GIST single-cell transcriptomics

The in-house ZZU-GIST single-cell transcriptomic cohort was generated using GEXSCOPE microfluidic chips for single-cell barcoding, mRNA capture, cDNA synthesis, amplification, and library construction. For the in-house cohort and the two public GIST single-cell cohorts integrated into dbGIST, cells were retained after quality-control filtering based on detected gene number, total UMI count, mitochondrial transcript fraction, and doublet detection where applicable.

Expression matrices were normalized using the LogNormalize method, highly variable genes were selected for dimensionality reduction, and UMAP embeddings were generated from scaled principal-component inputs. Batch-effect assessment was performed by comparing analyses with and without Harmony integration. Before-QC, after-QC, and Harmony-comparison panels for all three scRNA-seq cohorts are shown in Supplementary Figure S4A–L, with detailed preprocessing procedures provided in Supplementary Methods Sections 1.4.4 and 2.4.

### 4.4 Online website and AI-assisted implementation

dbGIST is implemented as a modular web resource in which a React and TypeScript front end communicates through a Nginx reverse proxy with a Node.js and Express service layer and multiple independent R Shiny analysis modules. Omics-specific functions are organized as separate yet methodologically aligned applications for genomics, transcriptomics, proteomics, phosphoproteomics, single-cell transcriptomics, and non-coding RNA analysis. The platform also includes an optional AI-assisted interpretation layer that provides narrative explanation and guided query support without altering the underlying statistical results. External literature-retrieval resources, including OpenScholar, are provided only as optional user-support tools for gene-centric literature exploration. They were not used to generate, modify, or validate the omics analyses, statistical tests, survival analysis, enrichment analysis, immune deconvolution, drug-response prediction, experimental validation, or figures reported in this study. All numerical results were generated by predefined dbGIST analytical modules. In other words, the LLM component augments interpretation but does not modify the computational backend. Platform architecture, reproducibility details, and software-version records are summarized in Supplementary Methods Sections 3.1–3.3 and Supplementary Table S6.

### 4.5 Module-level analyses

Across transcriptomic, proteomic, and phosphoproteomic modules, dbGIST supports cohort-level clinical comparisons, correlation analyses, survival analyses, pathway interpretation, and response-associated modeling using the measurement scale appropriate for each omics layer. Group comparisons are visualized with boxplots or violin plots and tested with paired or non-parametric procedures according to study design and group structure. Survival associations are evaluated by Kaplan-Meier analysis with cutpoint-based dichotomization and log-rank testing, and drug-response associations are summarized either by ROC analysis where labeled response data exist or by pharmacogenomic prediction models where direct response labels are unavailable. Functional interpretation is based on over-representation analysis and gene set enrichment analysis, while immune-context analyses use established deconvolution frameworks applied within cohorts rather than on a pooled matrix. Full statistical settings are summarized in Supplementary Methods Section 2.

The non-coding RNA module is designed as a read-only evidence-aggregation layer that links user-specified genes to curated miRNA, circRNA, and lncRNA resources with evidence grading and source attribution. Together, these design choices keep the web platform computationally lightweight while preserving methodological transparency and reproducibility. Additional platform-level implementation and supplementary methodological detail are described in Supplementary Methods Sections 2 and 3.

### 4.6 Cell culture and transfection

The human gastrointestinal stromal tumor (GIST) cell lines GIST-T1 and GIST-882 were cultured in their respective recommended media supplemented with 10% fetal bovine serum (FBS). Cells were maintained at 37 °C in a humidified atmosphere containing 5% CO2. For gene silencing experiments, cells were transfected with specific small interfering RNAs (siRNAs) using Lipofectamine 2000 reagent (Life Technologies, USA), strictly adhering to the manufacturer’s protocol. Non-targeting siRNAs served as negative controls. All siRNAs were designed and synthesized by Sangon Biotech (China).

### 4.7 Cell viability assay (CCK-8)

Cell viability was determined using the Cell Counting Kit-8 (CCK-8) assay. Briefly, GIST-T1 and GIST-882 cells, subjected to the respective treatments, were seeded into 96-well plates at a density of 2 × 10³ cells per well. At 1, 2, 3, and 4 days after treatment, CCK-8 reagent was added to each well according to the kit’s instructions, followed by incubation for a specified duration. The absorbance of each well at 450 nm (OD450) was then measured using a microplate reader (DG5032, Hua Dong, Nanjing, China).

### 4.8 Wound healing assay

The migratory capacity of GIST cells was assessed using a wound healing assay. Transfected GIST-T1 and GIST-882 cells were plated in 6-well plates at a density of 1 × 10⁵ cells per well and allowed to grow until reaching over 90% confluence. A uniform wound was created in the monolayer by scraping with a sterile 200 μL pipette tip. The initial wound width (0 hour) was immediately recorded by capturing images under a microscope. The wound width was measured again after 48 hours of incubation to quantify cell migration.

### 4.9 Quantitative Real-Time PCR (qRT-PCR)

Total RNA was isolated from cultured cells using Tri-Reagent (TIANGEN BIOTECH, Beijing, China) following the supplier’s protocol. RNA concentration and purity were assessed spectrophotometrically. For each sample, 3 μg of total RNA was reverse-transcribed into complementary DNA (cDNA). Quantitative PCR was performed to amplify the genes of interest, with β-actin serving as an internal control for data normalization.

### 4.10 Western Blot Analysis

Cellular proteins were extracted using RIPA lysis buffer. Protein concentration was quantified with a BCA Protein Assay Kit. Equal amounts of protein (30 μg per lane) were separated by SDS-polyacrylamide gel electrophoresis (SDS-PAGE) and subsequently transferred onto polyvinylidene fluoride (PVDF) membranes. After blocking with 5% bovine serum albumin (BSA) in TBST for 1 hour at room temperature, the membranes were incubated overnight at 4 °C with primary antibodies against MCM7 (1:1000 dilution) and GAPDH (1:1000 dilution). Following extensive washing, the membranes were incubated with appropriate horseradish peroxidase (HRP)-conjugated secondary antibodies at 37 °C for 1 hour. Protein bands were visualized using an enhanced chemiluminescence (ECL-plus) detection kit, and band intensities were analyzed.

## 5. CONCLUSION

In conclusion, dbGIST establishes a GIST-focused multi-omics resource that integrates public and in-house cohorts with diagnostic verification, modality-aware preprocessing, curated clinical annotations, cohort-level analytical modules, and transparent supplementary source data. The platform supports both interactive web-based exploration and programmatic reuse through public APIs that expose selected derived analysis results, while the optional LLM-assisted layer improves navigation and interpretation without modifying the statistical backend. The MCM7 case study demonstrates how dbGIST can connect cross-dataset evidence, multi-omics context, and experimental follow-up to generate testable target hypotheses. By making GIST multi-omics evidence more traceable, reusable, and interoperable, dbGIST provides a practical resource for target exploration and precision oncology research in GIST.

## DATA AVAILABILITY

dbGIST is available as an online resource through https://www.dbgist.com. Dataset-level accession information, cohort manifests, clinical dictionaries, inclusion/exclusion records, and supplementary methodological details are provided in the Supplementary Methods and Supplementary Tables. Quantitative outputs underlying key figures are provided as source-data tables where applicable. Public datasets used in this study are accessible from GEO (https://www.ncbi.nlm.nih.gov/geo), ArrayExpress (https://www.ebi.ac.uk/biostudies/arrayexpress), GSA-Human/NGDC (https://ngdc.cncb.ac.cn/gsa-human/), iProX (https://www.iprox.cn), and cBioPortal (https://www.cbioportal.org/). Key code components are hosted on GitHub (https://github.com/FuLab-ZhaoSun/dbGIST). dbGIST also provides public API access to selected analysis results and the corresponding visualization-ready data, enabling external tools and workflows to programmatically query derived resource outputs. In-house proteomic and single-cell transcriptomic data were generated under institutional ethics approval 2024-KY-1592-001 and 2024-KY-2222-02 and with written informed consent.

## ACKNOWLEDGEMENTS

This study was supported by Chinese National Natural Science Foundation (92474115 and 82261138558), the Joint Fund of Henan Provincial Research and Development Program for Science and Technology (242301420008), the Scientific Research and Innovation Team of the First Affiliated Hospital of Zhengzhou University (QNCXTD2023022), the Excellent Youth Fund of Henan Natural Science Foundation (212300410075), Zhengzhou University Young Student Basic Research Projects (PhD students) (ZDBJ202518).

## AUTHOR CONTRIBUTIONS

YF, JY, SL, and CL contributed to the conception and design of the study. ZS, QZ, JH-L, and JJ-L organized the database. ZS, FY, YT, XL, and YW developed the website and performed the statistical analyses. QZ, HL, and YG performed laboratory experiments. ZS, QZ, JH-L, and JJ-L wrote the first draft of the manuscript. SP, GZ, ZZ, MY, WM, and GL wrote sections of the manuscript. All authors contributed to manuscript revision, read, and approved the submitted version.

## COMPETING INTERESTS

The authors declare no competing interests.

## Notes

### Competing Interest Statement

The authors have declared no competing interest.

